# “GPress: a framework for querying General Feature Format (GFF) files and feature expression files in a compressed form”

**DOI:** 10.1101/833087

**Authors:** Qingxi Meng, Idoia Ochoa, Mikel Hernaez

## Abstract

1

**Motivation:** Sequencing data are often summarized at different annotation levels for further analysis. The general feature format (GFF) and its descendants, the gene transfer format (GTF) and GFF3, are the most commonly used data formats for genomic annotations. These files are extensively updated, queried and shared, and hence as the number of generated GFF files increases, efficient data storage and retrieval are becoming increasingly important. Existing GFF utilities for accessing these files, like *gffutils* and *gffread*, do not focus on reducing the storage space, significantly increasing it in some cases. Hence, we propose *GPress*, a framework for querying GFF files in a compressed form. In addition, GPress can also incorporate and compress feature expression files, supporting simultaneous queries on both files.

**Results:** We tested GPress on several GFF files of different organisms, and showed that it achieves on average a 98% reduction in size, while being able to retrieve all annotations for a given identifier or a range of coordinates in a few seconds. For example, on a Human GFF file, GPress can find all items with a unique identifier in 2.47 seconds and all items with coordinates within the range of 1,000 to 100,000 in 4.61 seconds. In contrast, gffutils provides faster retrieval but doubles the size of the GFF files. When additionally linking an expression file, we show that GPress can reduce the size of the expression file by more than 92%, while still retrieving the information within seconds. GPress is freely available at https://github.com/qm2/gpress.

## 2 Introduction

Uncovering the genetic structures and variations of the genome, as well as understanding their biological meaning, is one of the final goals of next-generation sequencing data analysis. To meet this goal, genomic sequences are often summarized at different annotation levels. The general feature format (GFF) and its descendants, such as the gene transfer format (GTF) and GFF3, are among the most commonly used data formats to provide genomic annotation information for next-generation sequencing data analysis [1]. In particular, the GFF files are used for describing genes and other features of DNA, RNA and protein sequences, and they contain several annotations for each feature, such as location and attributes. Given its importance in fundamental biological research, the GFF format is commonly used in many large genomic projects, such as the 5000 Insect Genomes Initiative (i5k) [2], the Plant Genome Initiative [3] and the Genome 10K (G10K) [4]. Although the number (and size) of GFF files may still be of lesser concern than sequencing files (such as FASTQ and SAM/BAM files), GFF files are more frequently revised, annotated, queried and streamed. For example, the i5k hosts more than 55 insect genome projects and uses the manual annotation program Apollo [5] to perform community annotation. Hence, for each project, community annotation can generate thousands of updates or revisions [6]. As such, methods to reduce the storage footprint of these data while providing efficient access and update capabilities are needed.

However, currently available GFF utilities, such as *gffread* [7] and *gffutils* [8], only aim at processing or accessing the data efficiently, and do not focus on minimizing the required space. In particular, gffread is used to filter, convert or cluster GFF records, but works directly on the original file; gffutils, on the other hand, imports the GFF file into a local sqlite3 file-based database, allowing users to search and access the data efficiently from it. However, the created database is generally larger than the size of the input file. Hence, we developed *GPress*, a framework for efficiently querying compressed GFF files (GFF3 and GTF files). In particular, GPress supports retrieving all features (and the corresponding fields) related to a given identifier, or that fall within a given range of coordinates. Further, to demonstrate the high modularity and flexibility of GPress, we included support for compressing and linking a feature expression file to the GFF file, allowing for simultaneously querying information from both files.

When tested on several GFF files, GPress is able to compress two times better than gzip on average and supports queries within seconds, while gffutils creates a database of doubled size on average.

## 3 Methods and experimental results

The encoder of GPress takes advantage of the correlations between different columns in the GFF file, and applies transformations to them for improved compression (see Supplementary data). Streams of data of similar characteristics are then grouped together and compressed with the general lossless compressor BSC [9]. GPress supports selective access over the compressed data (that is, random access), retrieving all information related to a set of identifiers or a range of coordinates. To achieve this, GPress compresses the data in blocks, and creates several index tables that record the structure of the data for fast retrieval, and hence it prevents unnecessary decompression. Lastly, a file containing expression values (in the form of, for example, counts, tpm, and/or fpkm) together with additional annotations such as estimated lengths for the different features can be integrated into the compressed GFF file. Moreover, GPress links the new information to the compressed GFF file, since items from the GFF file and the expression file may share the same unique identifier, and supports simultaneous selective retrieval from both files (see Supplementary data).

For the analysis, we selected three GFF files, hereafter denoted as *file 1, 2*, and *3*, of *H. Sapiens, M. Musculus*, and *D. Rerio*, which occupy 1.14GB, 0.8GB, and 0.43GB, respectively. These files serve as a representative of the many annotation files that exist for their same species (as the differences between GFF files from the same species are minimal), and hence they reflect the characteristics of a larger group of GFF files. To asses the performance of GPress integrating expression files into compressed GFF files, we used an expression file derived from the raw sequencing data of the TCGA-BRCA project and linked it to the Human GFF file (see Supplementary Data). All experiments were performed on an Intel i7 64-bit laptop with 16GB of RAM and 450GB of disk memory. GPress uses one thread, and in all conducted experiments the memory usage was less than 30MB for compression and 10MB for decompression.

We first analyze the performance of GPress when using a single block during compression. Note that in this case random access is not supported, as the whole compressed file needs to be decompressed to access the original data. GPress achieves an average compression ratio of 64.56, which is about 2.5 times higher than that of gzip (Table 1). In terms of running time, GPress is slower but comparable to gzip (Table 1).

**Table 1:** Performance of GPress without random access, and comparison to gzip when applied directly to the original file. The compression ratio (CR) is computed as “original size / compressed size”. CS, CT and DT stand for compressed size, compression time, and decompression time, respectively.

|  | CS [MB] |  | CR |  | CT [sec] |  | DT [sec] |  |
| --- | --- | --- | --- | --- | --- | --- | --- | --- |
|  | gzip | GPress | gzip | GPress | gzip | GPress | gzip | GPress |
| file 1 | 41.30 | 17.30 | 27.60 | 65.90 | 15.15 | 36.17 | 8.07 | 42.56 |
| file 2 | 28.10 | 11.90 | 28.47 | 67.23 | 9.91 | 27.98 | 5.58 | 24.84 |
| file 3 | 17.60 | 7.10 | 24.43 | 60.56 | 6.33 | 13.87 | 2.60 | 12.92 |

Next, we assess the performance of GPress when random access is enabled. Due to space constraints, here we provide results when considering blocks containing the information of 2,000 genes, as it provides a good trade-off between compression efficiency and time retrieval, and refer to the Supplementary data for results on other block sizes. We report retrieval time (averaged over ten queries) for queries based on an identifier and a range of coordinates. The range size varies from 1 × 10^4^ to 1 × 10^8^ (see Supplementary data for the effect of the range size). For a fair comparison, gzip is run on each of the data stream generated by GPress, for each block. To resolve queries based on an identifier, blocks are decompressed sequentially until the sought elements are found, whereas for queries based on range, we do a binary search. Searches on the original (uncompressed) file are treated similarly. GPress offers the best trade-off in compression ratio and retrieval time, achieving a compression ratio about two times higher than gzip while significantly reducing the search time (Fig. 1). On the other hand, gffutils is the fastest on retrieval, but at the cost of creating a database that doubles the size of the original file (Fig. 1). As an example, when tested on *file 1*, GPress can find all items on chromosome 1 with coordinates within the range of 1,000 to 100,000 in 4.61 seconds, and all items with unique identifier “ENSE00001612954.1” in 2.47 seconds. See Supplementary data for detailed results.

**Figure 1:**
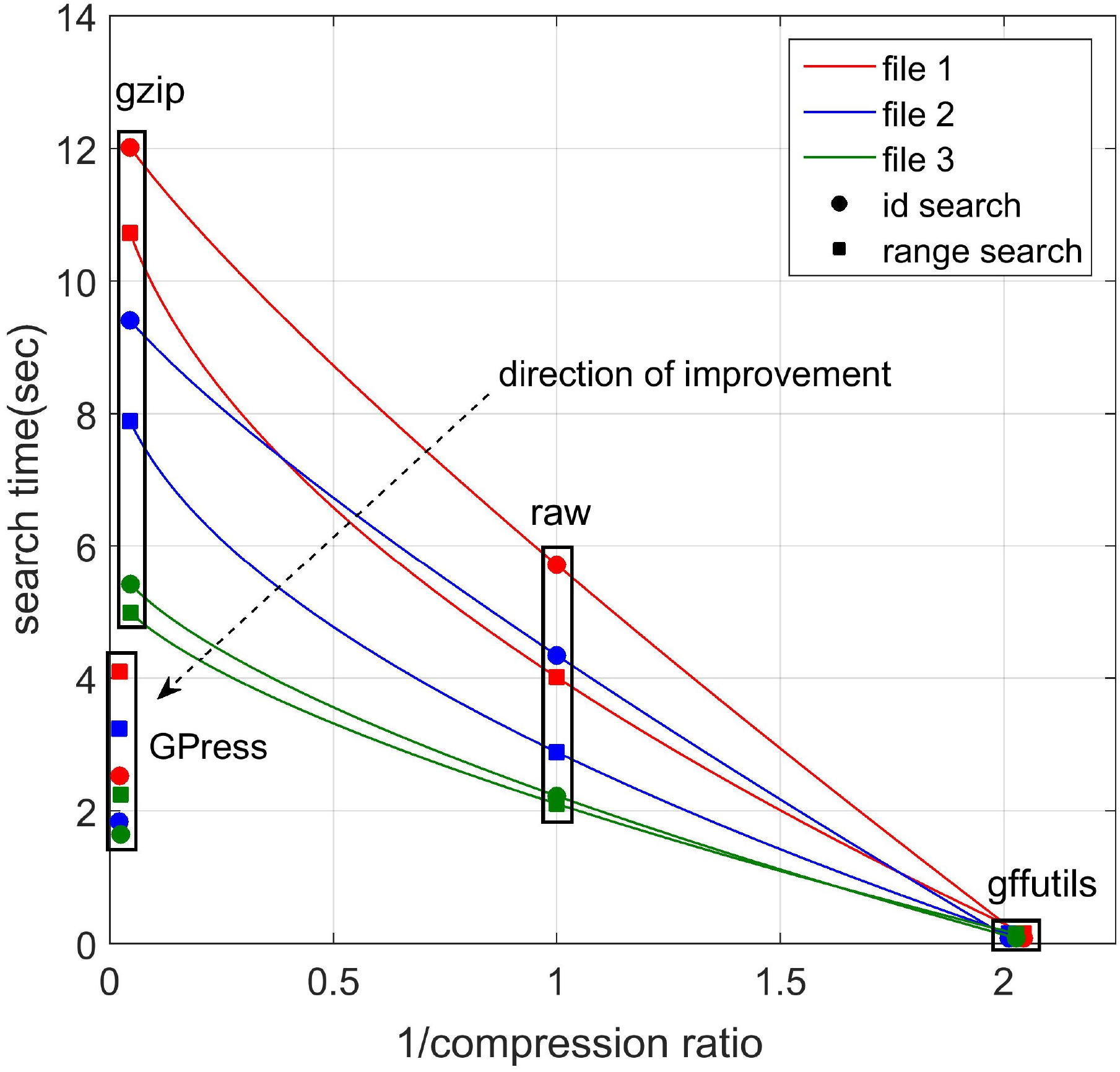
Comparison of search time and CR of the original file, GPress, gzip, and gffutils.

Finally, we analyze the performance when the expression file is compressed and linked to the compressed *file 1*. GPress reduces the size of the file from 7.7GB to 0.6GB, in less than 7 minutes. In addition, queries based on an identifier and on a range of coordinates are resolved in less than 3 and 5 seconds, respectively (averaged over 10 queries). Note that elements contained in both the GFF and the matrix file are retrieved. These results demonstrate the potential of GPress to compress and link additional files, while being able to retrieve information in seconds. In summary, GPress offers a framework with high modularity and flexibility that could be extended to support additional genome annotation information.

## 4 Method: GPress

### 4.1 GFF/GTF files

The general feature format (GFF) is a data format heavily used for storing annotation information. These include the gene transfer format (GTF) and the GFF3, which are currently the most commonly used versions. These files have one line per annotation, and each line consists of 9 columns (or fields), tab-separated. In particular, each column contains the following information:

- column 1 (string): *seqname*, the name of the sequence.
- column 2 (string): *source*, a unique label indicating the name of the program that generated this feature, or the data source.
- column 3 (string): *feature*, the feature type name (e.g., “gene” or “exon”).
- column 4 (integer): *start*, the start position of the feature in the sequence (sequence numbering starting at 1).
- column 5 (integer): *end*, the end position of the feature in the sequence (sequence numbering starting at 1).
- column 6 (floating point): *score*, a degree of confidence in the feature’s existence and coordinates.
- column 7 (character): *strand*, the strand of the feature, defined as “+” (forward) or “-” (reverse).
- column 8 (integer): *frame*, phase of the CDS features; it can be one of 0, 1, 2.
- column 9 (string): *attributes*, a semicolon-separated list of tag-value pairs, providing additional information about the feature.

The annotations of the GFF files are sorted by *seqname* (column 1), and within a given sequence (or chromosome), by the starting position of the genes, in increasing order. Each gene annotation is then followed by all annotations belonging to that gene. Within each gene, the columns are sorted by the starting position of transcripts in increasing order. Similarly, within each transcript, the exons and other items are sorted by the starting position. The order is increasing on a sense strand or decreasing on an antisense strand.

### 4.2 GPress encoder

During compression, GPress reads the GFF file line by line, and generates different streams of data for efficient compression. In particular, due to the different nature of each column, GPress parses them differently. In the following we describe this process in more detail.

The seqname (column 1), source (column 2), feature (column 3) and attributes (column 9) consist of strings. The values of these columns tend to be the same for consecutive records, and thus they are relatively easy to compress. Therefore, GPress writes them directly into 4 separate streams.

The start (column 4) and end (column 5) consist of integer numbers. Since the difference between end and start is much smaller than the end itself, GPress keeps the start and the difference between the start and the end. In order to further make the integers smaller and easier to compress, GPress keeps the difference between the next feature’s start and the current feature’s start or end. For features like transcripts that could overlap with each other, GPress keeps the difference between the next feature’s start and the current feature’s start. For other features that do not overlap with each other, GPress keeps the difference between the next feature’s start and the current feature’s end, so as to make the difference as small as possible. Hence, GPress parses the starting positions in the following way. If a gene is the first one in the file, GPress keeps its start unmodified. Otherwise, GPress keeps the difference between its start and the previous gene’s end. If a transcript is the first one in a gene, GPress keeps the difference between its end and the previous gene’s start. Otherwise, GPress keeps the difference between its end and its previous transcript’s start. GPress applies the same rule for exons and other features. In order to avoid negative number, GPress makes sure the difference is always non-negative by checking the strands. Finally, GPress writes the difference between the current end and the current start in a stream, and the difference between the current start and the previous end or start into a separate stream.

The score (column 6) could be an integer or a floating number. However, all the exons and introns are annotated based on alignments of RNA-seq reads, cDNAs and/or ESTs, and can be considered real, so filtering them based on a score is generally not considered. Therefore, those features usually do not have scores. At the transcript level, the transcripts are scored according to how well mRNA and EST alignments match over its full length. However, these scores depend on the experimental data and vary depending on the source. Therefore, GPress does not modify the scores (column 6) and write them directly into a stream.

The strand (column 7) indicates the sense strand of the feature and it can take values “+” (positive, or 5’->3’), “-”, (negative, or 3’->5’), and “.” (undetermined). Within a gene, all the items (transcripts, exons, etc.) have the same strand value. Therefore, GPress only keeps the strand value of the genes and writes them into a stream. All other strand values are discarded since they can be recovered given the strand value of the genes. The frame (column 8) indicates the phase of the features associated to the CDS feature type and can be 0, 1 or 2. Since only the CDS, the start codon and the stop codon belong to the CDS feature type, features with other feature types (e.g., transcripts, exons, etc.) do not have frame values. For all those cases, the frame value is just “.” (not available). Therefore, the frame (column 8) is highly correlated with the feature (column 3). Furthermore, we observed that most of the items with the same feature type are likely to have the same frame value. For example, most of the start codons in a file may have the same frame value 1. Therefore, GPress generates three different streams of frame values, one for the CDS’s frame values, one for the start codon’s frame values, and one for the stop codon’s frame values. Since each of those three stream tends to have the same frame value, it is relatively easy to compress them. GPress discards all other features’ frame values because their frame values are “.” and thus can be easily recovered.

Once every column in the GFF file has been parsed and written into 11 separate streams, GPress compresses each of the streams with BSC. BSC is a high performance compressor based on lossless, block-sorting data compression algorithms. There are several advantages to BSC. First of all, BSC supports multiple algorithms that allows software fine-tuning for maximum speed or compression efficiency. We tuned the parameters of BSC for each of the generated streams, to maximize compression efficiency. Second, BSC uses in-place compression and decompression to save memory. Finally, BSC supports GPU acceleration using NVIDIA CUDA technology. Figure 2 provides a schematic of the compression scheme described above.

**Figure 2:**
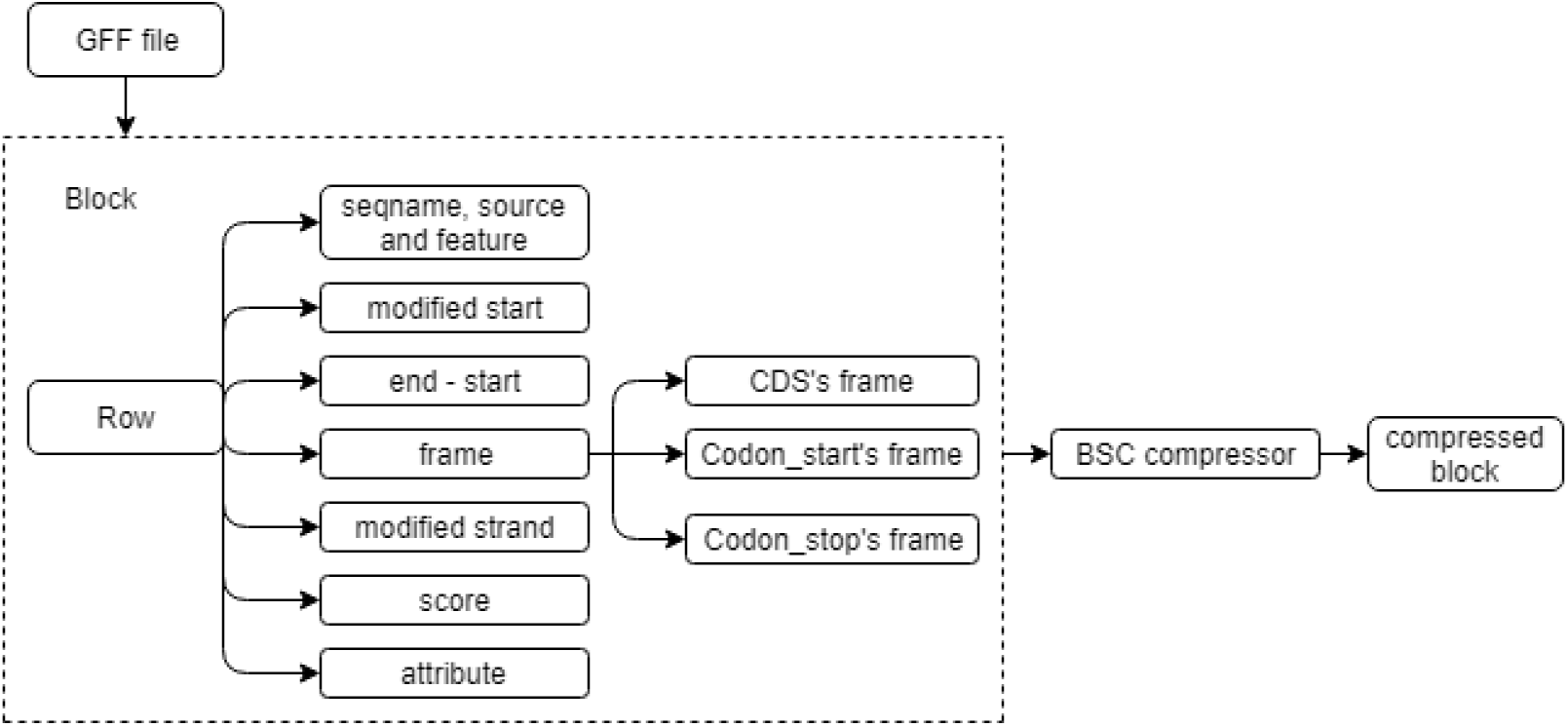
Schematic of the GFF compressor on a block extracted from the GFF file. GPress can be applied considering the whole GFF file as a unique block, but in that case selective access over the compressed data is not supported.

### 4.3 GPress decoder

During decompression, GPress first uses BSC to decompress all the streams generated during the encoding phase. GPress then reads each of the 11 streams line by line and reassembles them to reconstruct the original file. Specifically, GPress first reads in the feature (column 3) and strand (column 7). Then, GPress uses the feature and strand to recover the start (column 4), end (column 5) and frame (column 8), which are a mirror image processing with respect to the encoder. Finally, GPress reads in all the other columns and combines all the fields into one line (in the correct order) of the output file. GPress terminates once an EOF symbol is decoded from all streams at same time. If one file reaches the EOF before the other files, GPress produces an error message.

### 4.4 Random access (or selective access) over the compressed data

In addition to compression, GPress supports random access (or selective access) over the compressed data. To achieve this, the GFF file is first divided into blocks, and then each block is compressed separately. In this way, the access to an item or a group of items is more efficient since GPress only needs to decompress the blocks containing the sought items instead of the whole file. The compression method employed for each block is the same as the one described above, and blocks are constructed such that they contain all items belonging to a given number of genes (a user-selected parameter, 2,000 by default). Note that the resulting blocks may be of varying size (i.e., the number of items per block is not constant).

GPress supports two different types of queries (searches), one based on the unique identifier of an item, and the other based on a range of coordinates. If a unique identifier is provided, all items related to that identifier are retrieved. Similarly, given a range of coordinates, GPress retrieves all items that fall within that range. In order to determine which blocks need to be decompressed for a given query, GPress generates index tables during compression to map each item to a certain block. Specifically, GPress uses a hashtable to map each unique identifier of an item into its block number and position within the block. The unique identifier in our implementation is the ID specified in the attribute (column 9). If the GFF file is formatted in a canonical way, the selected unique identifiers work well to identify each item uniquely. In addition, GPress uses an index table to keep the starting and ending block numbers for each seqname (column 1). Because the coordinates of the items are only unique within each sequence name in a GFF file, the query on coordinates must also specify the interested sequence name. Thus, the index table of the starting and ending block number for each sequence name can help GPress to pinpoint relevant blocks quickly. Finally, GPress also constructs a hashtable to keep the starting and ending coordinates of each block. The starting coordinate of a block is the starting position of the first item in the block. The ending coordinate of a block is the ending position of the last item in the block. This index table is very helpful for querying on a range of coordinates because GPress can skip blocks that have starting or ending coordinates outside of the interested range. The schematic of generating the index tables is shown in Figure 3.

**Figure 3:**
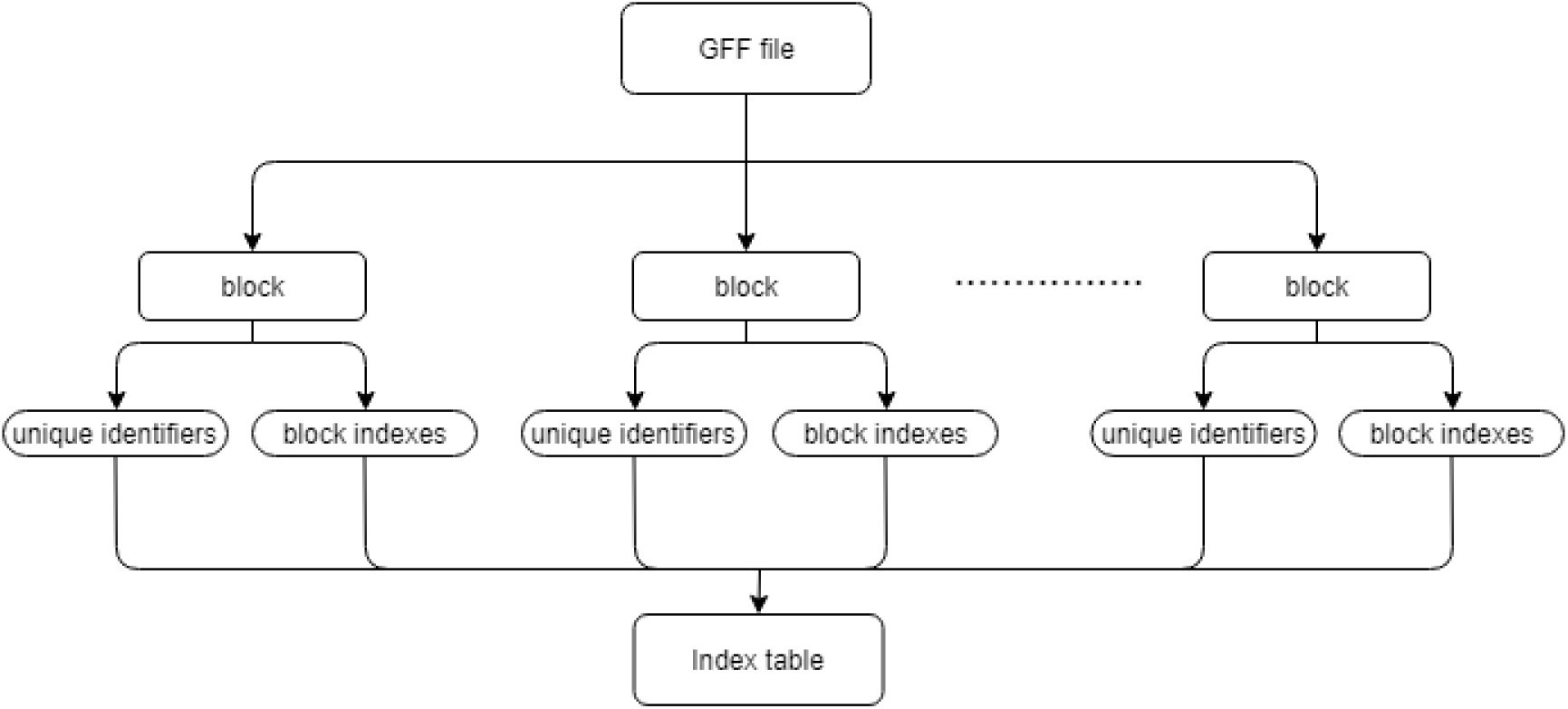
Schematic of producing the index tables.

Given a query, GPress therefore performs the following operations. When a unique identifier is provided, GPress uses the index tables to find the block number that contains the items related to that identifier, and the specific position within the block. Then, GPress decompresses that block in the same way as that stated in the decompression section. Since GPress also knows the specific position of the item within the block, it can easily find all the information of the items related to the searched item and retrieve it (e.g., if the unique identifier of a gene is provided, GPress will retrieve all the items pertaining to that gene, including transcripts, exomes, etc.). GPress can also search for all the items within a range of coordinates in a certain sequence name provided a range of coordinates and a sequence name. GPress pinpoints the relevant blocks for the sequence name of interest by using the index tables. Then, GPress checks the starting and ending coordinates of each of the relevant blocks to further rule out irrelevant blocks. GPress then decompresses the relevant blocks and retrieves all items that have coordinates within the specified range.

### 4.5 Expression files

GPress also provides the framework to compress and link the information contained in a expression file to that of a compressed GFF file. Expression files contain gene or transcript expression values, and they can be sparse or non-sparse. Here we focus on the non-sparse expression files, generally resulting from bulk RNA-seq analyses. These files usually consist of 6 columns, each column containing the following information:

- column 1 (string): *target ID*, unique identifiers within different databases
- column 2 (string): *sample*, the sample number
- column 3 (floating point): *EST counts*, counts of expressed sequence tags
- column 4 (floating point): *tpm value*, number of transcripts per million
- column 5 (floating point): *effective length*, the length of transcript after correction
- column 6 (integer): *length*, the regular length of the transcript

GPress reads in the non-sparse expression file line by line. First, GPress checks the target ID (column 1) of the item. The target ID consists of unique identifiers of the item within one or more databases, and includes identifiers for both transcripts and genes. GPress first checks whether or not one of the listed unique identifiers in the target ID already exists in the compressed GFF file by checking the index table of GFF’s unique identifiers. If the item contains a unique identifier that already exists in the previously compressed GFF file, GPress adds all other new unique identifiers to the GFF’s index table which all point to the same item. Otherwise, GPress adds all the unique identifiers in the existed hashtable and points them to a new item. Then, GPress writes the sample (column 2), EST counts (column 3), tpm values (column 4), effective length (column 5) and length (column 6) into five separate data streams. Once all lines of the expression file are read, the data streams are further divided in blocks and compressed with BSC (block size is defined as before, with 2,000 genes per block by default). In addition, to support random access, index tables are created in the same way as for the GFF file.

The information in the expression file can therefore be efficiently recovered given a unique identifier, in a similar fashion as for the case of GFF files. In particular, given a unique identifier contained in the expression file, GPress will retrieve all information related to it from the expression file, as well as from the GFF file (if available). Similarly, if a unique identifier from the GFF file is provided, GPress will retrieve all items from the GFF file related to it, as well as from the expression file (if available). With respect to queries based on a range of coordinates, note that the expression files do not contain this information. Therefore, queries based on a range of coordinates will first retrieve all relevant items from the GFF file. Then, given the unique identifiers from those items, GPress will retrieve all information related to those identifiers from the expression file (using the index tables).

In summary, given a unique identifier within one or more databases, GPress can find and retrieve all its related information in both the GFF file and the expression file. If a range of coordinates for a given sequence name is provided instead, all relevant items from the GFF will be retrieved, as well as any additional information from those items contained in the expression file. The schematic of the compression for an expression file is shown in Figure 4.

**Figure 4:**
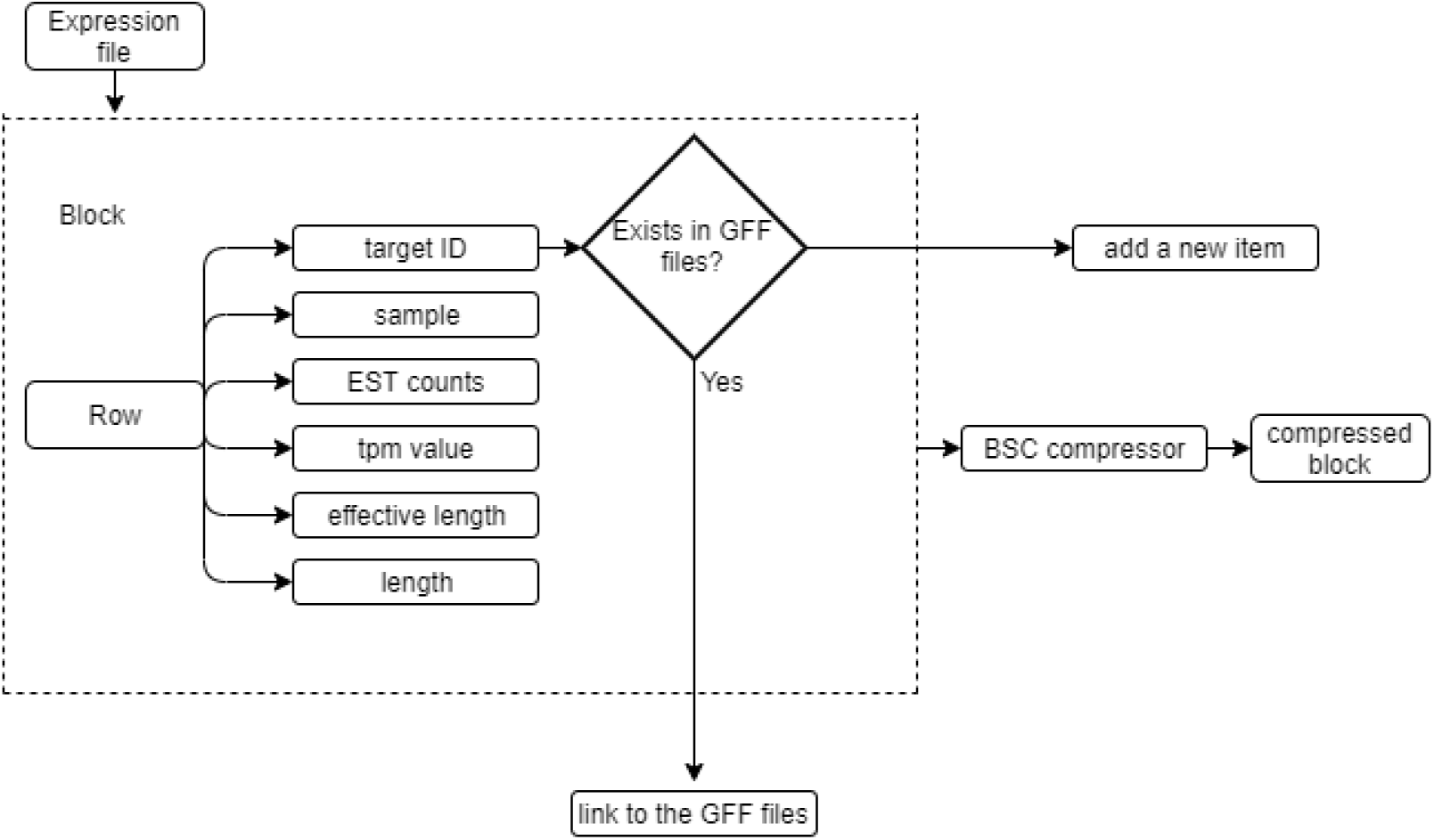
Schematic to link and compress an expression file in the GPress framework.

## 5 Results

To assess the performance of GPress, we used the data files specified in Table 2. These files include three GTF files from different organisms and an expression file. The file number specified in Table 2 is coherently used in both the main paper and the supplementary (i.e., the main paper refers to these files by their file number as defined in the table). The expression matrix file (file 4) was generated from the raw sequencing data of the TCGA-BRCA project, and hence we just used a generic name for it.

**Table 2:** Description of the files used in the experiments.

| Number | File names | File size [GB] | Source | Description |
| --- | --- | --- | --- | --- |
| 1 | *gencode.v30.annotation.gtf* | 1.14 | GENCODE | Comprehensive gene annotation for *Homo sapiens* |
| 2 | *gencode.vM21.annotation.gtf* | 0.80 | GENCODE | Comprehensive gene annotation for *Mus musculus* |
| 3 | *Danio_rerio.GRCz11.96.gtf* | 0.43 | GENCODE | Comprehensive gene annotation for *Danio rerio* |
| 4 | *expresion_file.tsv* | 7.73 | TCGA-BRCA | Expression file for *Homo sapiens* |

The performance of GPress when no random access is supported (i.e., the entire GFF file is considered as a single block during compression) is reported in the main paper, and hence in the following we focus only on the performance of GPress with random access (or selective access) over the compressed data. Performance in this case is measured in terms of compression ratio and execution time of queries.

Tables 3, 4 and 5 summarize the performance of GPress when applied to files 1, 2, and 3, respectively, for different block sizes (recall that the results presented in the main paper correspond to 2,000 genes per block). As expected, increasing the number of genes per block reduces the total number of blocks. The total compression size reported in the tables includes the size of the compressed GTF file as well as the size of the index tables. The compression ratio represents the ratio between the original size and the total compression size achieved by GPress. The compression time includes the time to compress the files and to generate the index tables, and the loading time indicates the time needed to load the tables prior to performing queries. Finally, the *id* and *range* searches stand for the average search time given a unique identifier or a range of coordinates, respectively. The reported time for the id search is the average over ten queries with different unique identifiers and the reported time for the range search is the average of ten queries with range size varying from 1 × 10^4^ to 1 × 10^8^.

**Table 3:** Performance of GPress with different block sizes on file 1, *gencode.v30.annotation.gtf*.

| block size<br>[gene] | num. of<br>blocks | size index<br>tables<br>[MB] | total<br>compression<br>size [MB] | compression<br>ratio | compression<br>time[sec] | loading<br>time [sec] | id search<br>[sec] | range<br>search [sec] |
| --- | --- | --- | --- | --- | --- | --- | --- | --- |
| 500 | 116 | 7.36 | 26.86 | 42.44 | 123.28 | 3.38 | 1.45 | 7.84 |
| 1000 | 57 | 7.39 | 25.89 | 44.03 | 79.24 | 3.36 | 1.82 | 5.83 |
| 1500 | 38 | 7.39 | 25.79 | 44.20 | 63.43 | 3.39 | 2.02 | 5.43 |
| 2000 | 28 | 7.38 | 25.58 | 44.57 | 58.66 | 3.41 | 2.53 | 4.56 |
| 2500 | 23 | 7.39 | 25.39 | 44.90 | 52.36 | 3.39 | 3.55 | 4.86 |
| 3000 | 19 | 7.41 | 25.21 | 45.22 | 48.82 | 3.42 | 3.85 | 5.33 |
| 3500 | 16 | 7.44 | 25.14 | 45.35 | 46.75 | 3.43 | 4.12 | 5.67 |

**Table 4:**
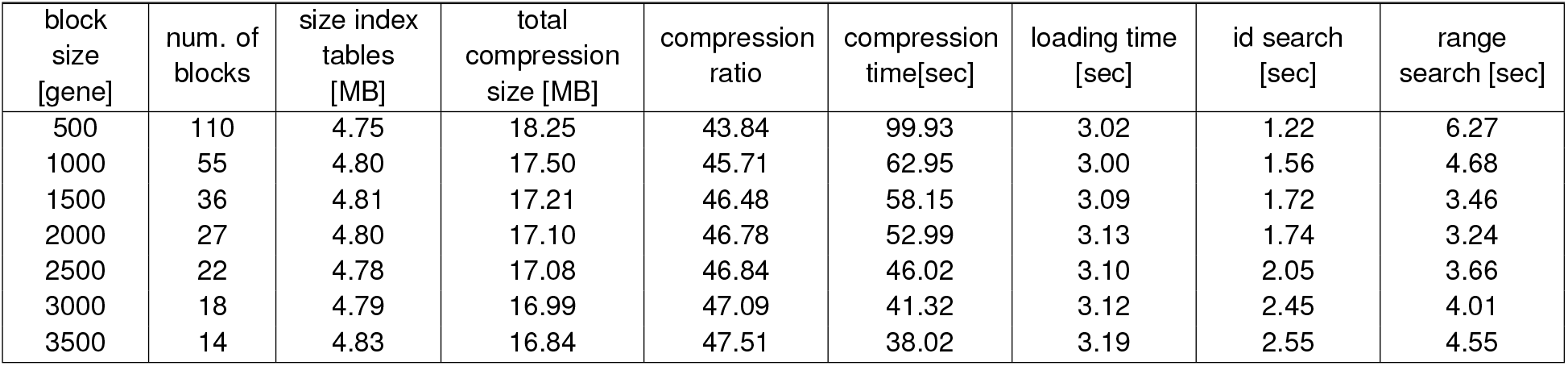
Performance of GPress with different block sizes on file 2, *gencode.vM21.annotation.gtf*.

| block<br>size<br>[gene] | num. of<br>blocks | size index<br>tables<br>[MB] | total<br>compression<br>size [MB] | compression<br>ratio | compression<br>time[sec] | loading time<br>[sec] | id search<br>[sec] | range<br>search [sec] |
| --- | --- | --- | --- | --- | --- | --- | --- | --- |
| 500 | 110 | 4.75 | 18.25 | 43.84 | 99.93 | 3.02 | 1.22 | 6.27 |
| 1000 | 55 | 4.80 | 17.50 | 45.71 | 62.95 | 3.00 | 1.56 | 4.68 |
| 1500 | 36 | 4.81 | 17.21 | 46.48 | 58.15 | 3.09 | 1.72 | 3.46 |
| 2000 | 27 | 4.80 | 17.10 | 46.78 | 52.99 | 3.13 | 1.74 | 3.24 |
| 2500 | 22 | 4.78 | 17.08 | 46.84 | 46.02 | 3.10 | 2.05 | 3.66 |
| 3000 | 18 | 4.79 | 16.99 | 47.09 | 41.32 | 3.12 | 2.45 | 4.01 |
| 3500 | 14 | 4.83 | 16.84 | 47.51 | 38.02 | 3.19 | 2.55 | 4.55 |

**Table 5:** Performance of GPress with different block sizes on file 3, *Danio_rerio.GRCz11.96.gtf*.

| block size [gene] | num. of blocks | size index tables [MB] | total compression size [MB] | compression ratio | compression time[sec] | loading time [sec] | id search [sec] | range search [sec] |
| --- | --- | --- | --- | --- | --- | --- | --- | --- |
| 500 | 63 | 2.64 | 11.32 | 37.99 | 53.24 | 1.34 | 1.01 | 3.21 |
| 1000 | 31 | 2.67 | 10.84 | 39.67 | 43.12 | 1.31 | 1.38 | 2.69 |
| 1500 | 20 | 2.70 | 10.62 | 40.49 | 33.40 | 1.32 | 1.63 | 2.32 |
| 2000 | 15 | 2.72 | 10.44 | 41.19 | 28.52 | 1.25 | 1.69 | 2.25 |
| 2500 | 11 | 2.72 | 10.10 | 42.57 | 25.92 | 1.40 | 1.83 | 2.31 |
| 3000 | 9 | 2.73 | 10.04 | 42.83 | 20.99 | 1.39 | 2.14 | 2.34 |
| 3500 | 7 | 2.75 | 9.98 | 43.09 | 18.23 | 1.49 | 2.25 | 2.58 |

From the results in Tables 3, 4 and 5, we can see that the compression ratio increases as the block size increases. This is reasonable as larger block sizes group more similar data in one block, making them easier to compress. Similarly, the average id search time increases as the block size increases. This is also reasonable, since larger block sizes make GPress spend more time on decompressing each block. The average range search time first increases and then decreases as the block size increases. This may be caused by the fact that the range search time depends on both the size of each block and the number of decompressed blocks. When the block size is very small, the influence of the number of blocks dominates over that of each block size. However, when the block size becomes large, the size of each block becomes more important and dominates the number of decompressed blocks. Therefore, the best block size will depend on the application. For example, if most queries are id searches, a smaller block size is preferred. For a general application, a block size of medium length is the best as it offers a good trade-off between the id search and the range search, and the compression performance. By default, GPress operates with a block size of 2,000 genes.

For the default setting of 2,000 genes per block, Table 6 compares the performance of GPress with that of *gzip* and *gffutils*. These results are the ones used to generate Figure 1 of the main paper. As mentioned in the main paper, GPress offers better compression ratios than gzip and gffutils, while resolving the queries in a few seconds. In particular, the compression ratio is in general twice of that of gzip, and about an order magnitude higher than that of gffutils. In terms of retrieval time, gffutils is the fastest, employing less than a second on average, but GPress employs less than 3 seconds for the id queries and less than 5 seconds for the range queries.

**Table 6:** Performance of GPress with 2,000 genes per block and comparison to gzip and gffutils. CS, CR and CT stand for compression size, compression ratio, and compression time. The id and range search stand for the average search time in seconds given a unique identifier or a range of coordinates, respectively. STD corresponds to the standard deviation.

|  | CS [MB] |  |  | CR |  |  | CT [sec] |  |  |
| --- | --- | --- | --- | --- | --- | --- | --- | --- | --- |
|  | gzip | gffutils | GPress | gzip | gffutils | GPress | gzip | gffutils | GPress |
| file 1 | 52.30 | 2330 | 25.58 | 21.79 | 0.489 | 44.57 | 45.54 | 818.41 | 58.66 |
| file 2 | 36.61 | 1610 | 17.10 | 21.85 | 0.497 | 46.78 | 38.66 | 523.77 | 52.99 |
| file 3 | 20.11 | 872 | 10.44 | 21.38 | 0.493 | 41.19 | 22.72 | 283.13 | 28.52 |

Table 6: Performance of GPress with 2,000 genes per block and comparison to *gzip* and *gffutils*.
|  | id search (STD) [sec] |  |  | range search (STD) [sec] |  |  |
| --- | --- | --- | --- | --- | --- | --- |
|  | gzip | gffutils | GPress | gzip | gffutils | GPress |
| file 1 | 12.02 (5.32) | 0.0874 (0.0002) | 2.53 (0.062) | 10.73 (3.15) | 0.162 (0.0045) | 4.56 (0.350) |
| file 2 | 9.41(3.92) | 0.0871 (0.0002) | 1.74 (0.053) | 7.89 (2.73) | 0.158 (0.0043) | 3.24 (0.315) |
| file 3 | 5.12 (2.14) | 0.0848 (0.0002) | 1.69 (0.046) | 4.89(1.69) | 0.152 (0.0039) | 2.25 (0.262) |

Since for the range queries the number of decompressed blocks depends on the range size, next we analyze how the range search time varies with the size of the range. The results for all three considered files are summarized in Figure 5, for a block size of 2,000 genes. For each range size, the reported search time, in seconds, is calculated as the average over ten queries. The *extra small* corresponds to a range size around 1 × 10^4^, the *small* to a range size around 1 × 10^5^, the *medium* to a range size around 1 × 10^6^, the *large* to a range size around 1 × 10^7^, and the *extra large* to the range size around 1 × 10^8^. We observe that the range search time increases as the range size increases.

**Figure 5:**
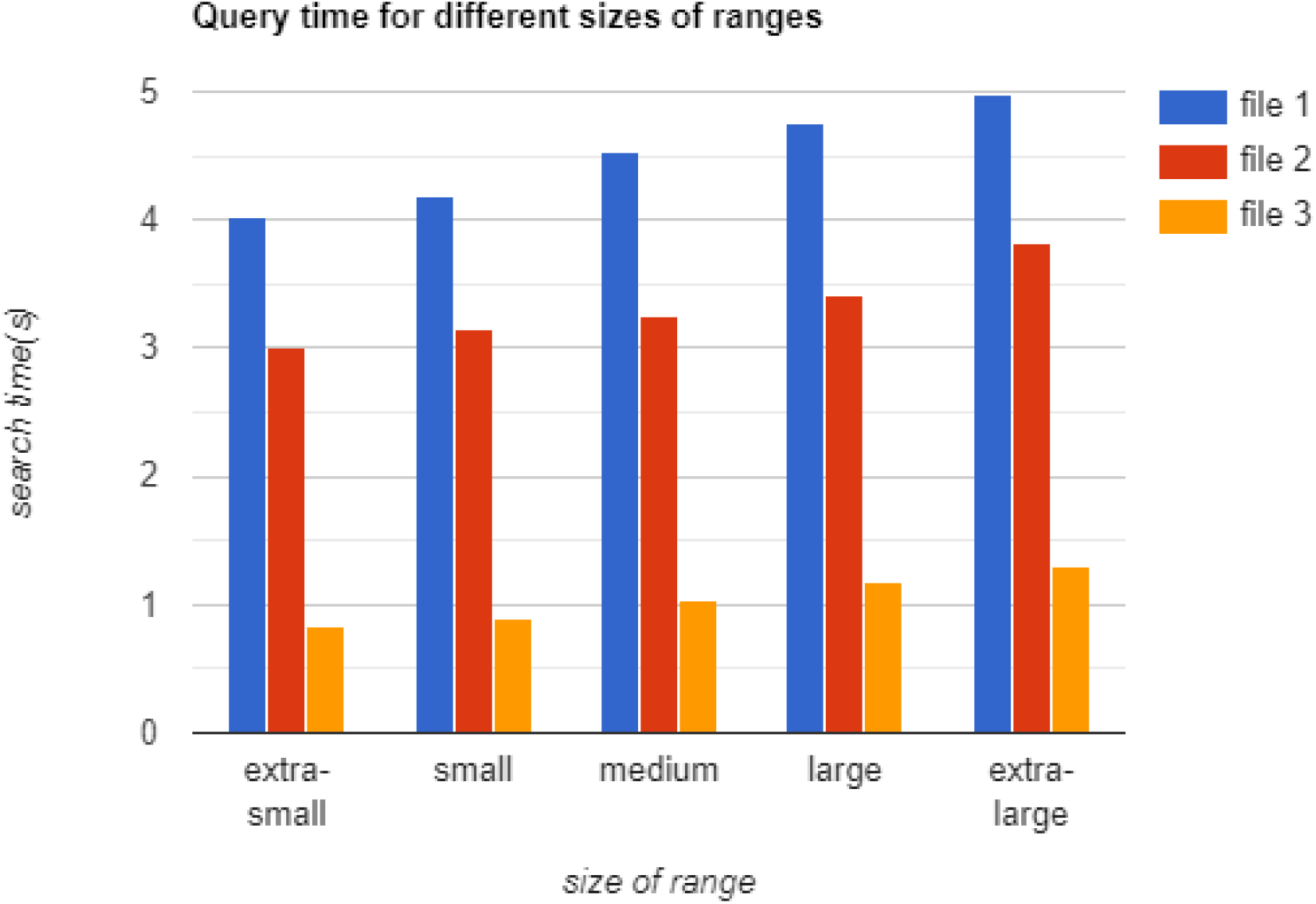
The retrieval time for range queries for different range sizes.

Finally, we analyze the performance of GPress when a expression file is compressed and linked to a compressed GTF file. In particular, we use the H. Sapiens annotation file (*file 1*) and the expression file (*file 4*) specified in Table 2. The results are summarized in Table 7. The original size refers only to the size of the expression file, and the same block size is used for both the GTF and the expression file. Similarly, the index table size, the total compression size, and the compression ratio refer only to the expression file. On the other hand, since the expression file contains items with identifiers that already exist in the GTF file, the id and range search referred to in the Table find relevant information in both the GTF and the expression files. From the results reported in the table, we observe that the influence of the block size on the expression file is nearly the same as on the GTF files, i.e., the compression rate decreases with the number of blocks. The compression rate is however smaller than that achieved on the GFF files. Regarding the retrieval time, we observe that there is less than a second increment with respect to querying only the GFF file, for both id and range searches. In summary, GPress is able to add new information to the framework without sacrificing on the query retrieval time.

**Table 7:** Performance of GPress when the expression file *(file 4)* is compressed and linked to *file 1*, for different block sizes.

| block size [gene] | number of blocks | original size [MB] | index table size [MB] | total compression size[MB] | compression ratio | id search [sec] | range search [sec] |
| --- | --- | --- | --- | --- | --- | --- | --- |
| 1500 | 50 | 7,730 | 1.99 | 596 | 12.97 | 2.39 | 5.52 |
| 2000 | 37 | 7,730 | 2.00 | 590 | 13.10 | 2.98 | 4.94 |
| 2500 | 30 | 7,730 | 2.00 | 587 | 13.17 | 4.01 | 5.12 |
| 3000 | 25 | 7,730 | 2.01 | 584 | 13.24 | 4.34 | 5.66 |
| 3500 | 21 | 7,730 | 2.01 | 570 | 13.56 | 4.57 | 5.91 |

## 6 Discussion

As the amount of GFF files increases, utilities that compress these files while supporting selective access (or random access) over the compressed data become imperative. Currently, GFF files are mainly compressed by means of the general lossless compressor *gzip*. Unfortunately, gzip performs poorly in compression, since it is not tailored to the data contained in GFF files. More importantly, gzip does not support random access. On the other hand, commonly available GFF utilities only aim at processing and accessing the data efficiently, regardless of the required space. For example, commonly used utility *gffutils* doubles the size of the original GFF file, as shown in Table 6. GPress, however, can reduce the size of the gzipped GFF files by half, while resolving queries in a few seconds. Supported queries include retrieving all items related to a unique identifier as well as all items that fall within a given range of coordinates. With respect to gffutils, the query time of gffutils is faster than that of GPress, but the compression ratio of GPress is around 90 times better than that of gffutils. As the number of generated GFF files that need to be queried increases, the advantage offered by GPress will become more apparent. In addition, GPress offers an added advantage, as it can also compress expression files and linked them together to the compressed GFF files. Queries in this case look for the relevant items in both the GFF and the expression files, and are still resolved in a few seconds. Finally, the framework of GPress could be extended in the future to support other annotation formats in addition to expression files. Since there is a trend in the field of genomics to develop methods that can compress arbitrary file formats, the modularity and flexibility of GPress are very beneficial in this case.

## Notes

https://github.com/qm2/gpress

## References

[1] H. Zhang, Overview of Sequence Data Formats. New York, NY: Springer New York, 2016, pp. 3–17. [Online]. Available: https://doi.org/10.1007/978-1-4939-3578-9_1

[2] J. D. Evans, “The i5k initiative: Advancing arthropod genomics for knowledge, human health, agriculture, and the environment.” Journal of Heredity, vol. 104, no. 5, pp. 595–600, 2013. [Online]. Available: http://www.library.illinois.edu.proxy2.library.illinois.edu/proxy/go.php?url=http://search.ebscohost.com.proxy2.library.illinois.edu/login.aspx?direct=true&db=asn&AN=89734146&site=eds-live&scope=site

[3] B. Jeffrey L., K. Elizabeth A., L. Michael, and M. Joachim, “A plant genome initiative.” The Plant Cell, vol. 10, no. 4, p. 488, 1998. [Online]. Available: http://www.library.illinois.edu.proxy2.library.illinois.edu/proxy/go.php?url=http://search.ebscohost.com.proxy2.library.illinois.edu/login.aspx?direct=true&db=edsjsr&AN=edsjsr.10.2307.3870727&site=eds-live&scope=site

[4] K.-P. Koepfli, B. Paten, and S. J. O’Brien, “The genome 10k project: a way forward.” Annual Review Of Animal Biosciences, vol. 3, pp. 57–111, 2015. [Online]. Available: http://www.library.illinois.edu.proxy2.library.illinois.edu/proxy/go.php?url=http://search.ebscohost.com.proxy2.library.illinois.edu/login.aspx?direct=true&db=cmedm&AN=25689317&site=eds-live&scope=site

[5] E. Lee, G. A. Helt, J. T. Reese, M. C. Munoz-Torres, C. P. Childers, R. M. Buels, L. Stein, I. H. Holmes, C. G. Elsik, and S. E. Lewis, “Web apollo: a web-based genomic annotation editing platform.” Genome Biology, vol. 14, no. 8, p. R93, 2013. [Online]. Available: http://www.library.illinois.edu.proxy2.library.illinois.edu/proxy/go.php?url=http://search.ebscohost.com.proxy2.library.illinois.edu/login.aspx?direct=true&db=cmedm&AN=24000942&site=eds-live&scope=site

[6] M.-J. M. Chen, H. Lin, L.-M. Chiang, C. P. Childers, and M. F. Poelchau, The GFF3toolkit: QC and Merge Pipeline for Genome Annotation. New York, NY: Springer New York, 2019, pp. 75–87. [Online]. Available: https://doi.org/10.1007/978-1-4939-8775-7_7

[7] G. Pertea, “gffread,” version 0.11.5. [Online]. Available: http://ccb.jhu.edu/software/stringtie/gff.shtml

[8] R. Dale, gffutils, version 0.8.4. [Online]. Available: https://pythonhosted.org/gffutils/

[9] I. Grebnov, bsc and libbsc, version 3.1.0. [Online]. Available: http://libbsc.com/

